# Generation of a Ym1 Deficient Mouse utilising CRISPR-Cas9 in CB6 Embryos

**DOI:** 10.1101/2025.03.10.642405

**Authors:** J E Parkinson, G E Baldwin, P H Papotto, N E Humphreys, A D Adamson, J E Allen, T E Sutherland

**Author notes:** Equal contribution (A D Adamson, J E Allen, T E Sutherland).

## Abstract

Chitinase-like proteins (CLPs) are of wide interest due to their significant roles during both biological and pathological processes. Human CLPs such as YKL-40 have been suggested as biomarkers of disease severity in many conditions. Murine CLPs include Brp39, Ym1, and Ym2 and these are similarly upregulated in multiple mouse models of pathology. Investigation of these molecules, particularly Ym1 and Ym2, is plagued by complexity in the genomic locus due to recent gene duplication events in the C57BL/6 strain. Using a novel CRISPR-Cas9 targeting approach involving CB6 mixed background embryos, we generated a Ym1 deficient mouse. Validation using flow cytometry, ELISA, and immunofluorescence confirmed no expression of mature Ym1 protein with no alteration in the expression of related chitinases/CLP genes including Chia and Chil4. This new transgenic mouse line will be key for investigating CLP functions and the genetic approach utilised may provide a useful strategy for other genes which show differences between inbred mouse strains.

## Introduction

Chitinase-like proteins (CLPs) are expressed in a range of organisms including mammals^1^, insects^2–4^, parasites^5,6^, and even plants^7–9^. They are part of the glycoside hydrolase 18 family and have evolved from gene duplication events of the active chitinases, their ancestral precursors^1,10^. Chitinases catalyse the break down chitin, the second most abundant natural polymer on earth after cellulose. Chitin is a structural and protective polymer found in Fungi and several phyla of Animalia; notably Nematoda and Arthropoda. Today humans are widely exposed to chitin from its use in industry, medicine, and biotechnology. Chitinases evolved not only to act as host defence molecules against chitin containing pathogens but also function in degrading the chitin we are exposed to in day-to-day life. Whilst some CLPs can still bind chitin, they have lost this chitin degrading ability^8^. Despite this, CLPs maintain an expression pattern broadly associated with conditions of type-2 inflammation such as nematode infection or asthma^8,11–15^.

The CLP family contains a diverse array of genes across many species and these molecules often show high sequence homology. The different members are thought to have evolved from birth and death evolution under strong purifying selection^16^. This has resulted in the independent evolution of species specific genes, with orthologs generally being more closely related than paralogs^1^. Humans have two main CLPs, YKL-40 (*CHI3L1*), and YKL-39 (*CHI3L2*). YKL-40 in particular has been suggested as a biomarker of disease severity in many conditions^17–21^. Notably increased YKL-40 protein levels have been reported in the serum in some cohorts people with asthma^12,22^. Despite these observations, and GWAS studies highlighting *CHI3L1* as an asthma risk gene^23^, the exact function of YKL-40 is still not well understood.

In comparison to humans, mice have three coding CLPs, Brp39 (*Chil1*), Ym1 (*Chil3*), and Ym2 (*Chil4*) and these have also been widely implicated in a plethora of diseases^24^. Ym1 and Ym2 in particular have a very high sequence homology at the exonic and protein level. Despite this they exhibit distinct expression patterns in the mouse^25^, highlighting potential differences in function. Both these CLPs are also under purifying selection, suggesting they have necessary functions in health and/or disease^1^. Despite the identification of these genes almost forty years ago^26^ the mechanisms by which they function are still poorly understood. Given that Ym1 and YKL-40 have been identified as highly upregulated proteins in many disease settings, the lack of in vivo tools to study them hinders understanding of their roles in disease pathogenesis.

The CRISPR-Cas9 editing system was initially discovered in bacteria and archaea where it functions to protect against invasive viral and plasmid nuclei acids^27,28^. Since then it has been adapted to work as a DNA editing system in eukaryotic cells^29–31^. CRISPR-Cas9 has now become the gold standard for the generation of genetically altered mice^32^. In this study we sought to generate a Ym1-deficient mouse strain. Due to genetic complexities this ultimately required the use of CRISPR-Cas9 mediated deletion in a hybrid background strain. The generation of this novel Ym1-deficient strain will enable detailed investigation of the biological functions of Ym1 and its role in disease aetiology and pathogenesis across many conditions.

## Results

### Chil3 and Chil4 encode for highly homologous CLPs

Across species CLPs show high sequence homology. Ym1 (*Chil3*) and Ym2 (*Chil4*) in particular are >90% homologous at the protein, RNA, and DNA level. Previous research has been hindered by the lack of tools available to accurately differentiate between the two molecules. This is despite several differences in the amino acid sequence, imparting distinct biochemical properties (**Fig 1a**). Both *Chil3* and *Chil4* have arisen from a recent gene duplication event, as evidenced by their location immediately adjacent on chromosome three (**Fig 1b**). Within the human and mouse chitinases and CLPs, *Chil3* and *Chil4* are unique as they show the highest sequence homology to each other, whereas most other CLPs and chitinases show greater homology to their orthologs, rather than associated paralogs. (**Fig 1c**). Recently tools for identifying the specific spatial distribution of these proteins have been developed demonstrating very different cellular expression patterns between Ym1 and Ym2 in the murine lung^25^ prompting further questions into their differing roles and how these align with human CLP orthologues.

**Figure 1:**
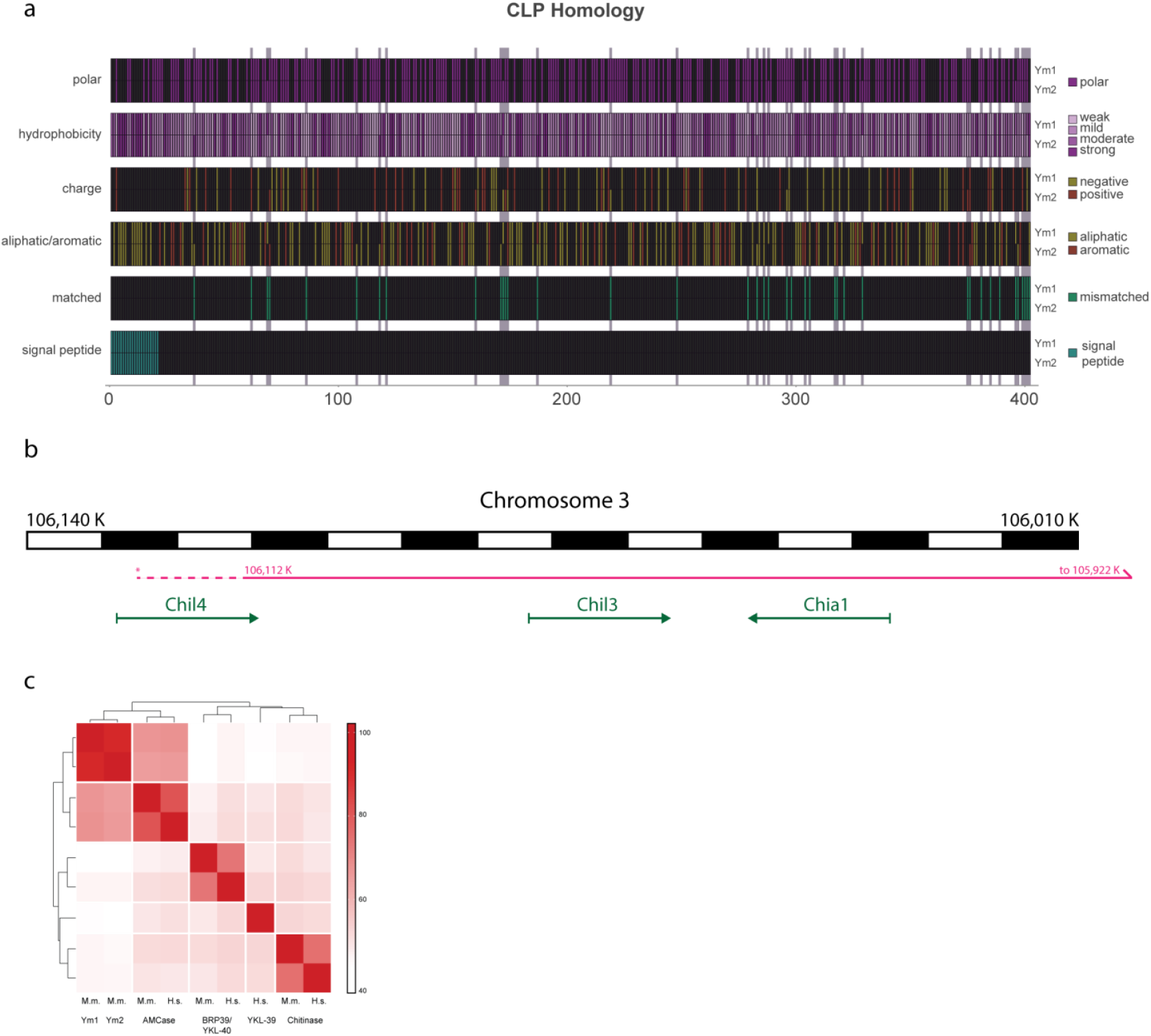
The two murine chitinase like proteins (CLPs) Ym1 and Ym2 have high sequence homology but have distinct differences at the protein sequence level. **a**) Schematic showing the differences in the amino acid sequence between the two proteins and associated differences in biochemical properties. The genes that encode Ym1 (*Chil3*) and Ym2 (*Chil4*) are also located immediately adjacent to each other on chromosome 3, **b**) Schematic showing the location of *Chil3* and *Chil4* and in relation to *Chia1* which encodes for AMCase. Black and white alternating blocks represent 10Kb of DNA, Solid magenta line shows the duplication suggested by Graubert *et al*^34^. Dashed magenta line shows an extension that is supported by the data within this manuscript. Relevant genes and their orientation on chromosome 3 are shown by green arrows. **c**) Heatmap showing the amino acid sequence similarity between the extended members of the chitinase and CLP family. Scale shows amino acid sequence similarity in percent. (M.m = Mus. musculus and H.s = Homo. Sapiens).

### Strategy for generating a Chil3 floxed mice

Gene loss through CRISPR-Cas9 mediated deletion is a classic approach to study gene function. However, the *Chil3* gene is particularly challenging to target due to its high sequence homology to *Chil4*^10^, found ∼30kb downstream on the same chromosome (**Fig 1b**). Whilst design of specific *Chil3* single guide RNAs (sgRNA) proved extremely difficult, we were able to exploit the intronic sequences which are sufficiently different between *Chil3* and *Chil4* to allow e (sg)-RNAs (g988 and g946) (**Fig 2a**). In addition to this approach, we planned to utilise a “floxed” allele strategy to allow spatiotemporal control of protein expression and to mitigate against potential lethality. We synthesised specific sgRNAs and cloned a complementary DNA repair template compromising a Homology-loxP-SP6R-Exon3-M13F-loxP-Homology sequence (**Fig 2a**). The sgRNA, Cas9 and deoxyribonucleic acid (DNA) donor were then co-delivered to C57BL6/J mouse embryos, which were transferred to pseudopregnant CD1 recipients.

**Figure 2:**
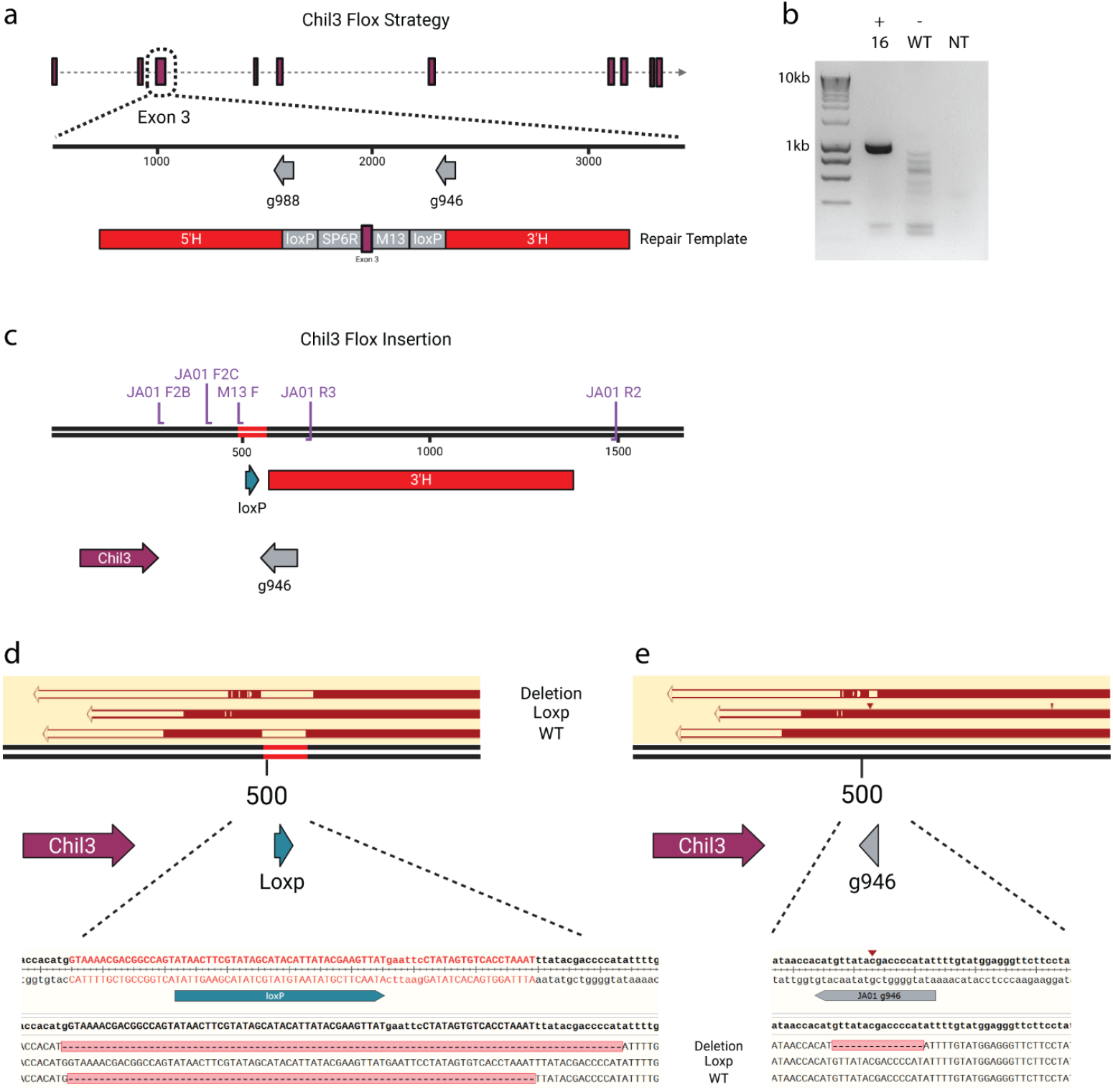
Genetic targeting strategy for generating a *Chil3* floxed mouse. **a**) Schematic showing the homology directed repair template including 3’ and 5’ homology arms and relative location relative to the *Chil3* gene. **b**) Genotyping results from the M13F and JA01 R2 primer sequences showing amplification of a 1kb product in pup 16 but not in C57BL/6 control DNA (WT) or non-template control (NT). **c**) Projected results from the methodology in a) and b) demonstrating the inclusion of an intronic LoxP site and flanking 3’ homology arm as well as respective primer binding sites. Sequencing results from pup 16 were transfected via blunt end ligation into plasmids and transformed colonies were miniprepped and Sanger sequenced using the M13R primer. (**d**-**e**) Sanger sequencing results of pup 16 aligned to the predicted HDR knock in sequence (**d**) and the Chil3 reference sequence (**e**) showing the presence of three different alleles including the LoxP insertion, 15bp deletion, and wild type sequence. Note the small red triangle indicates an insertion.

Screening of the initial F0 pups revealed low editing efficiency, but we obtained two mice with single LoxP integration, a common outcome of such experiments^33^. Pup 16 had integrated the 3’ LoxP site, as determined by PCR (**Fig 2b**) using the M13F primer sequence integrated as part of the donor template, and a Reverse primer targeted to the genome flanking the homology arm (JA01 R2) (**Fig 2c**). This LoxP site integration was then confirmed by Sanger sequencing of an amplicon generated by JA01 F1B and R2C (**Fig 2d and e**). Sequencing results revealed potential mosaicism as three alleles were detected: a WT allele, the LoxP allele, and a deletion of 15 bases (**Fig 2d and e**). An additional pup had integrated the 5’ LoxP, as determined using similar assays targeted to the 5’ end of the integration site (data not shown). We decided to establish a colony of mice bred from Pup 16 with a view to harvesting embryos from mice homozygous for the 3’ LoxP and eventually introduce the 5’ LoxP on this background using CRISPR and a single-stranded oligodeoxynucleotides repair template. To establish this colony, Pup 16 was crossed with a WT C57BL6/J male and germline transmission was confirmed using the same flanking PCR as before (**Fig 2c**). Then mice with the LoxP site (heterozygotes) were interbred to establish a homozygote colony.

### Identification of coinherited alleles and gene duplication of the CLP locus

We screened the F2 generation of pups using a PCR reaction (JA01 F1C/R3) that was designed to amplify smaller products of 136bp (WT) or 211bp (LoxP). This reaction allowed us to classify mice as being heterozygous or homozygous for the LoxP site more easily by size differences on a Qiaxcel automated gel system. However, we observed that the smaller amplicon, corresponding to the CRISPR induced deletion previously identified, routinely co-segregated with the LoxP site (**Fig 3a**).

**Figure 3:**
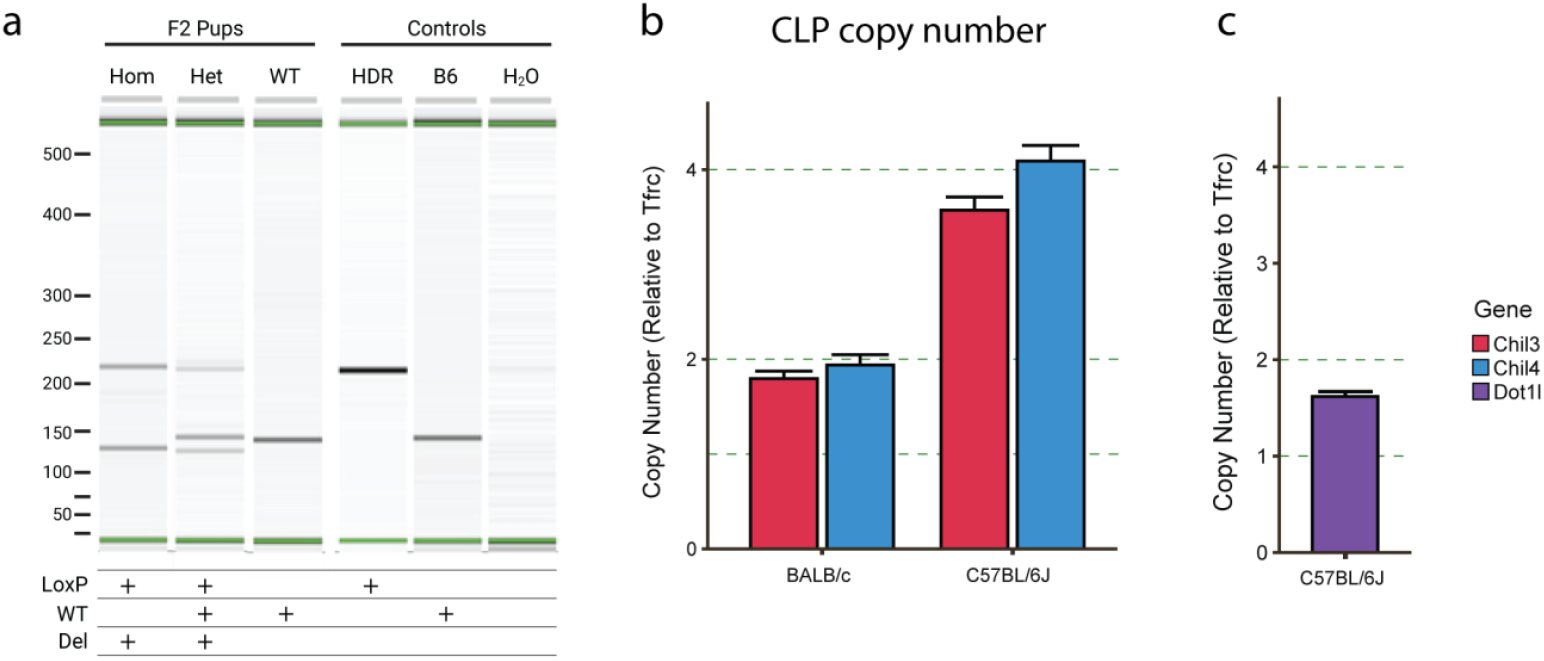
PCR Confirmation of the F2 generation from *Chil3* floxed mice. **a**) Genotyping results using a Qiaxcel automated gel system. Pups were genotyped for the presence of the LoxP or WT sequence using the (JA01 F1C/R3) primers. This showed the coinheritance of the LoxP site and the 15bp deletion that was observed previously as demonstrated in the homozygote mice which showed no presence of WT allele. WT mice showed no presence of the LoxP insert or the 15bp deletion and Het animals showed the presence of all three alleles. Bands were identified using controls including spiked HDR template (HDR) and WT C57BL/6J (B6) sequences in addition to a negative non-template control (H_2_O) shown on the right section of the plot. **b**) Confirmation of gene duplication using droplet digital PCR showing the copy number of *Chil3* and *Chil4* in BALB/c and C57BL/6J mice in reference to a control gene (*Tfrc*) with a copy number of two.

The co-segregation of both a Homology-directed repair (HDR) knock in and a non-homologous end joining (NHEJ) deletion in the same region led us to hypothesise that this region was duplicated in the C57BL/6 background strain. Using genomic DNA harvested from multiple mouse strains we performed ddPCR using a probe designed against *Chil3* and *Chil4* alongside a reference *Tfrc* probe. Analysis showed that there was a copy number duplication in the C57BL/6 strain but not in the BALB/c strain (**Fig 3b**). The copy number of the *Tfrc* gene was also cross-validated to another reference gene *Dot1l* (**Fig 3c**). The co-segregation of the two editing events can be explained by simultaneous targeting of the duplicated binding sites which exist on the same allele. After this observation, we noted that CLP locus duplication events had been reported previously in inbred mouse strains including C57BL/6 substrains, but notably not in other strains including BALB/c^34,35^. The presence of this recent gene duplication leads to complications with CRISPR targeting in the C57BL/6 strain.

### Utilising mixed background CB6 mice to generate a Chil3 deficient mouse

The CLP locus duplication in C57BL/6J led us to try an alternative knockout strategy, whereby we would delete the entire *Chil3* gene in an alternative mouse strain. BALB/c mice are a viable model for immunology research and lack the CLP duplication event according to ddPCR analysis (**Fig 3b**). Using BALB/c mice enables leverage of the intergenic regions for specific sgRNA design, whilst also allowing design of a single-stranded oligodeoxynucleotide DNA repair template for precise excision of the full gene (**Fig 4a**). However, standard superovulation and embryo harvesting techniques in BALB/c mice generated poor quality embryos with a low survival rate (64%) and few pregnancies when transferred to pseudopregnant females. Only 3 pups were born from a total of 346 embryos injected (0.9%) (**Table 1**). We reasoned that using CB6 (C57BL/6 x BALB/c) embryos would give us a *Chil3* allele to target from the BALB/c background, along with better embryo characteristics associated with the C57BL/6 strain. Indeed, the embryo retrieval and survival rate were improved compared to pure BALB/c embryos (81%). However, only a modest increase in pup numbers was observed, with 3 pups born from a total of 177 embryos injected (1.7%) (**Table 1**).

**Figure 4:**
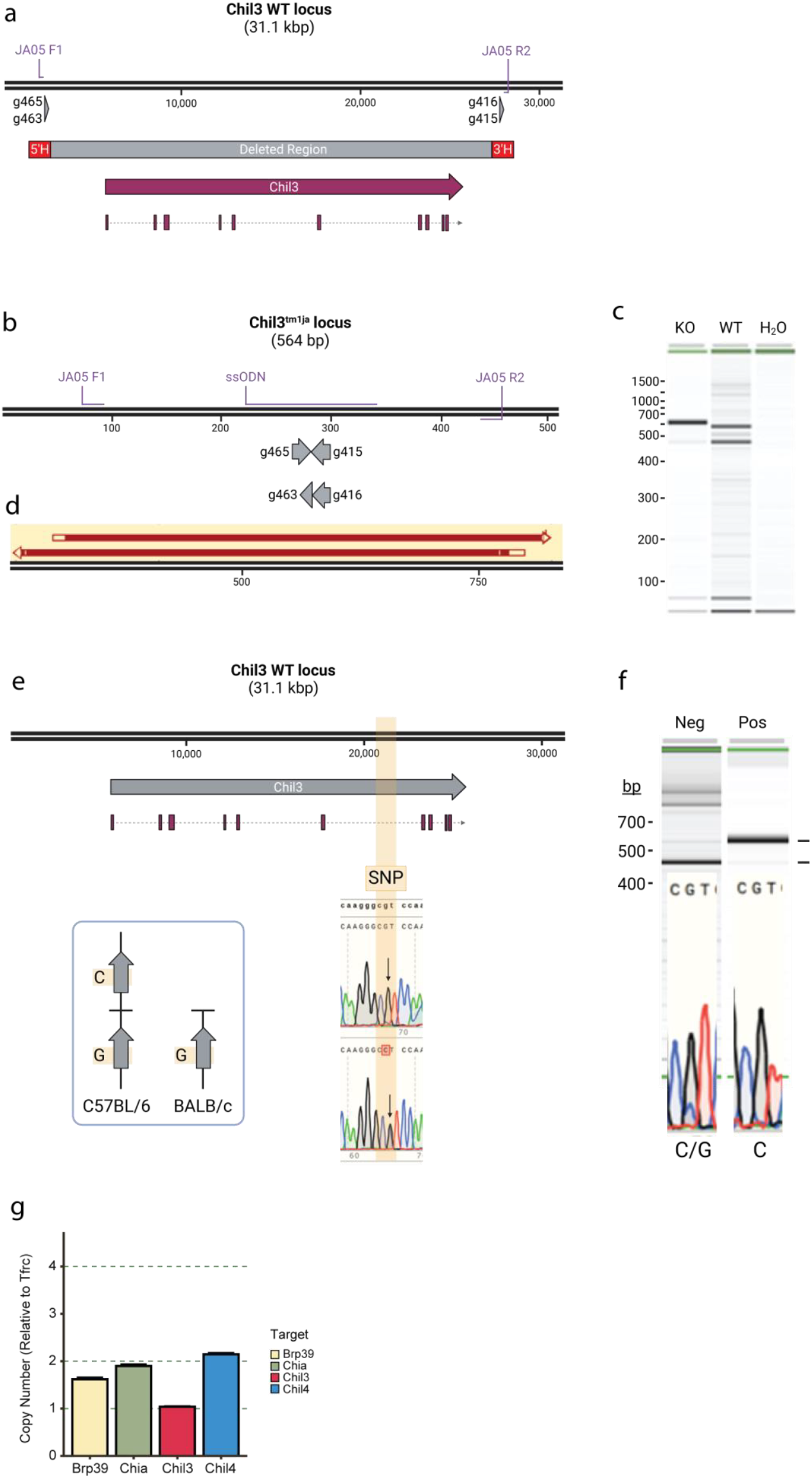
Strategy for generating a *Chil3*-deficient mouse. **a**) Schematic showing the guide RNAs and the expected deletion used in BALB/c and CB6 embryos. **b**) Schematic showing the expected deletion and respective oligo binding locations. **c)** PCR results using JA05 F1 and JA05 R1 amplifying a 564bp band in the KO animals, but not in the WT controls. **d**) Sanger sequencing confirming the sequence of the 564bp product in c) aligned with the predicted KO sequence in b). **e**) Schematic showing the location of strain specific SNPs in the *Chil3* gene with sanger sequencing confirming the sequence of these SNPs between C57BL/6 and BALB/c mice and the differences between gene copies in B6 animals. **f**) Sanger sequencing showing the presence of a pure C read in the SNP of a pup containing the deletion shown in a) and a 2:1 mixed G/C read in a littermate control without the mutation. **g**) Droplet digital PCR showing the copy number of *Chia1, Chil1, Chil3*, and *Chil4* in heterozygous Chil3-deficient offspring. Showing that there is a copy number of one for Chil3 showing that there has been a deletion of this gene, but with no change to the copy number of *Chil1, Chia1*, or *Chil4*.

The same CRISPR targeting was repeated in CB6 embryos in an attempt to specifically delete the BALB/c copy of *Chil3* (**Fig 4a**). Pups were genotyped using primers that would amplify deletion events (**Fig 4b**) but give no product in unedited mice (**Fig 4a, b and Table 2**). A single CB6 pup gave a band at the predicted size of 564bp (**Fig 4b, c and d**), which Sanger sequencing revealed to be a precise, HDR mediated deletion of the *Chil3* gene (**Fig 4d**). This founder was then backcrossed with a BALB/c mouse in order to confirm germline transmission of this deletion allele.

As the mutation could have targeted one of the duplicated C57BL/6 alleles, or the desired BALB/c allele, we needed to determine which copy of the *Chil3* gene was deleted in the founder. Potential single nucleotide polymorphisms (SNPs) between the strains that could be exploited to determine which allele had been deleted were selected from publicly available databases (http://www.informatics.jax.org/snp). This region of the mouse genome contained only one candidate SNP for this assay, an intronic base predicted to be a C nucleotide in C57BL/6 and a G nucleotide in BALB/c (**Fig 4e**). This sequence was amplified from both C57BL/6 and BALB/c genomic DNA and Sanger sequenced to confirm the webtool prediction. The BALB/c sequence was confirmed as a G nucleotide, but an overlapping peak indicated both a G and C nucleotide in the C57BL/6 strain (**Fig 4e**). Nevertheless, the SNP could still be exploited for genotyping (**Sup Fig 4a**). If the BALB/c *Chil3* allele was deleted in the founder, crossing with WT BALB/c mice would give a pure G read in this amplicon in F1 pups harbouring the deletion, whereas negative littermates without the deletion would transmit the C57BL/6 allele giving a mixed GC read. A pure G read was observed in F1 litters harbouring the deletion, indicating deletion of the desired BALB/c *Chil3* gene in the founder compared to control CB6 DNA which showed a mixed G/C read (**Fig 4e**). Final confirmation of appropriate copy number of *Chil3* was confirmed by ddPCR which showed that we could still detect the CLP duplication in C57BL/6 mice but this was not present in BALB/c controls or in the litter which contained the deletion, from the cross of a pure BALB/c and our founder animal (**Fig 4g**). In our animals we were also able to observe a loss of copy number between the BALB/c control and our F2 animals resulting from loss of the entire gene removing the binding sites for the primers and probes in our ddPCR assay (**Fig 4f and 3f**).

### Speed congenics allows rapid backcrossing of CB6 pups to common lab strains

There are large variances in responsiveness to certain pathogenic challenges between different inbred mouse strains^25,36^. To facilitate investigation into this and to provide a homogenous background on which to analyse these *Chil3*-deficient animals heterozygote animals were backcrossed to wildtype C57BL/6 and BALB/c animals. This allowed rapid stabilisation onto the BALB/c background using a speed congenic system (**Fig 5a**). Using a panel of strain specific SNPs (Transnetyx), the optimal mouse at each stage was use for generation of the next litter resulting in a fully backcrossed strain in five generations (**Fig 5b**).

**Figure 5:**
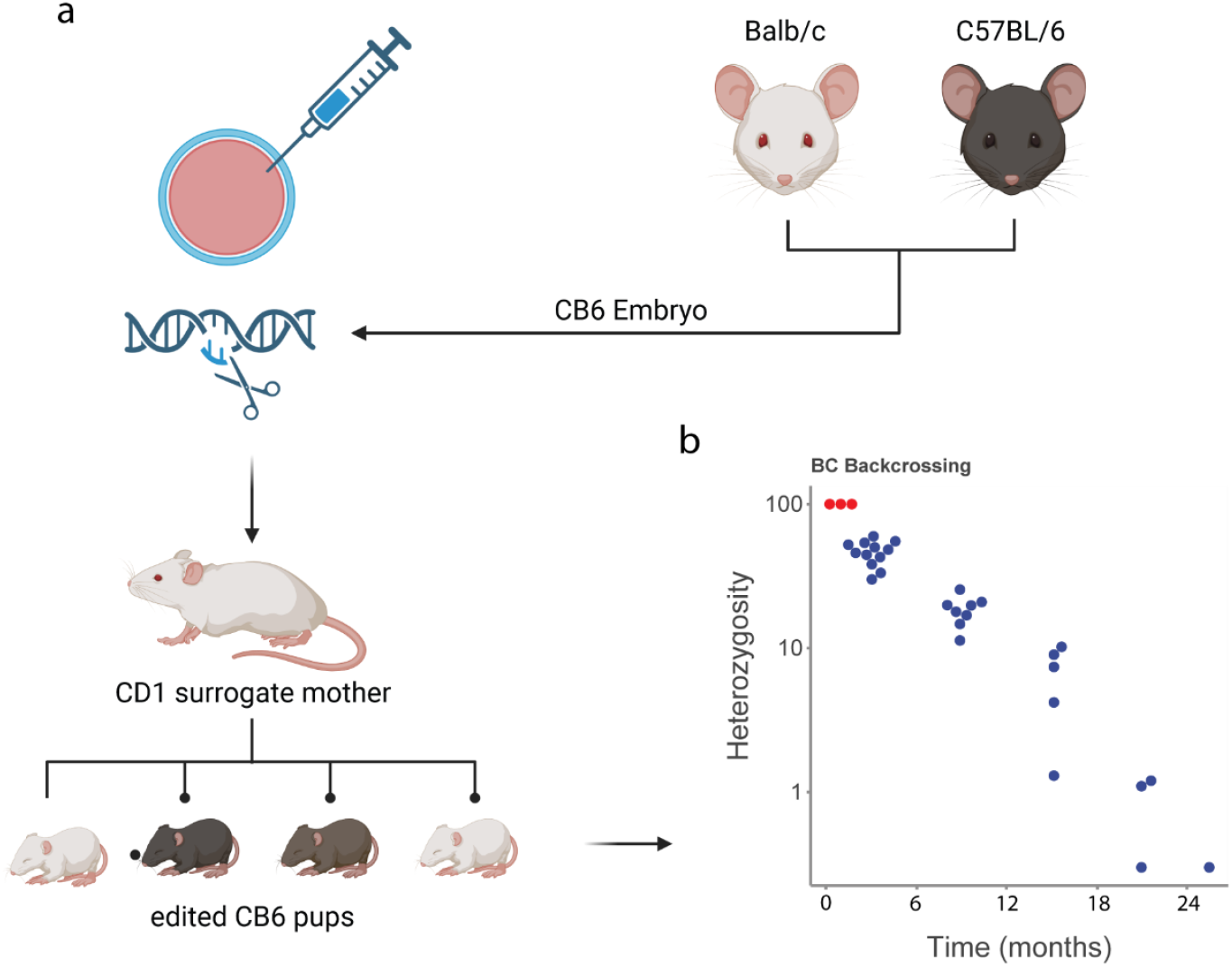
To facilitate scientific further characterisation and analysis the CB6 pups carrying the *Chil3* deletion were backcrossed onto a BALB/c background. **a**) strategy used to generate CB6 embryos for microinjection and **b**) subsequent heterozygosity of pups at each round of backcrossing showing completed transfer to the BALB/c background in five generations.

### Chil3-deficient animals lack of Ym1 protein with no change in related proteins

Ym1 protein, encoded by the *Chil3* gene, is abundant in murine lung in the steady state and known to be expressed by neutrophils and alveolar macrophages^37^. Therefore, we initially validated the loss of Ym1 in lung tissue from fully backcrossed BALB/c Ym1-deficient mice. We examined expression of *Chil3* by RT-qPCR in naïve mice and observed absence of *Chil3* mRNA in the lungs of knockout mice and a reduction of approximately half in heterozygous littermates (**Fig 6a**). This deficiency was also observed at a protein level in whole lung tissue homogenate (**Fig 6b**), as well as in the bronchoalveolar lavage (BAL) (**Fig 6c**). Again, heterozygous animals were found to have roughly half the level of Ym1 protein as wildtype littermates (**Fig 6b-c**). A similar reduction was observed in peritoneal fluid, although Ym1 expression in the peritoneal cavity was much lower than lung or BAL in the steady state (**Fig 6d**). Immunofluorescence staining of wildtype lung sections demonstrated Ym1^+^ cells throughout the lung parenchyma, consistent with its expression in alveolar macrophages^37,38^ (**Fig 6e**). Sections stained from Ym1-deficient mice showed no positive staining with heterozygote animals again demonstrating an intermediate protein expression level (**Fig 6e**), which was quantified at roughly half the expression level of Ym1 (**Fig 6f**). To confirm this quantitative difference the mean fluorescent intensity (MFI) of Ym1 between genotypes was measured by flow cytometry. A gradual reduction in MFI from wildtype to heterozygous to knockout animals was observed in histograms of lung alveolar macrophages (**Fig 6g and h**) and lung neutrophils (**Fig 6i and j**). There was no difference in pulmonary alveolar macrophage (**Fig 6k**) or neutrophils (**Fig 6l**) numbers between genotypes showing that the absence of Ym1 expression was not altering the number of these cell types in naïve mice.

**Figure 6:**
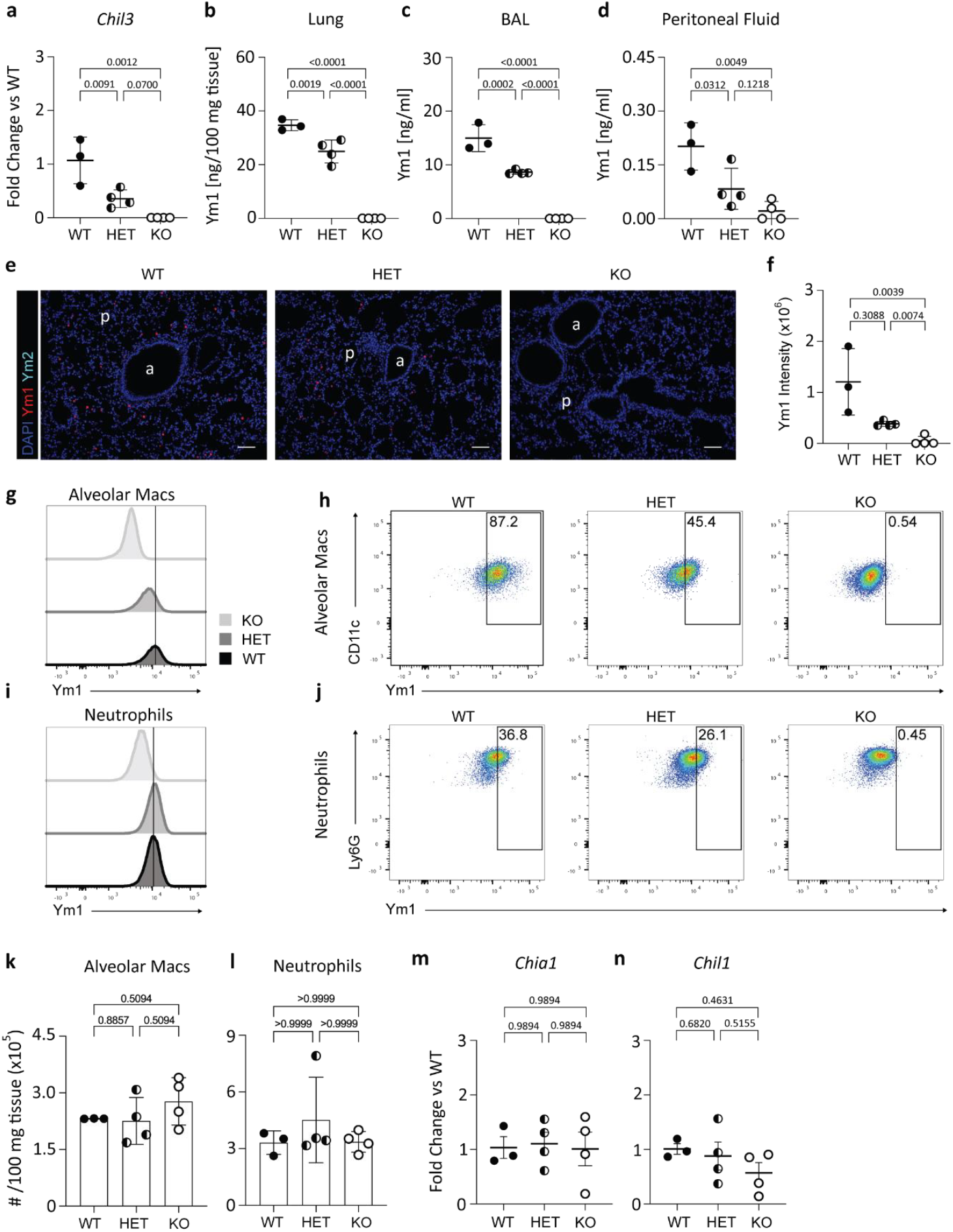
Phenotyping of the backcrossed BALB/c *Chil3*-deficient mice. (**a**) *Chil3* mRNA expression in wildtype (WT), heterozygous (HET), and knock out (KO) mice from whole lung tissue. (**b**) *Chil3* encodes the protein Ym1 which was quantified in whole lung homogenate normalised to the mass of the lung, (**c**) the bronchoalveolar lavage, and (**d**) peritoneal lavage from WT, HET, and KO Ym1*-*deficient mice. (**e**) Immunofluorescence showing the expression of Ym1 and Ym2 in lung tissue from naïve mice in WT, HET, and KO *Chil3* deficient mice. (**f**) Quantification of the fluorescent intensity of Ym1 in WT, HET and KO Ym1-deficient mice. Plots showing the MFI of Ym1 staining in alveolar macrophages (**g** and **h**) and Neutrophils (**i** and **j**) from WT, HET, and KO Ym1-deficient lungs relative to FMOs. (**k**) Total number of alveolar macrophages, and (**l**) neutrophils, identified from flow cytometry from digested whole lung tissue. RNA expression of active chitinase *Chia1* (**m**) and CLP *Chil1* (**n**) from whole lung in WT, HET, and KO Ym1-deficient mice. Data were analysed by ANOVA with Tukey’s multiple comparison test with significance level showing comparisons between each different genotype, p values are enumerated within each panel.

Due to the sequence similarity between family members of the chitinases and CLPs (**Fig 1**) we wanted to ensure that expression of these were not altered in our Ym1-deficient mice. *Chil4* (Ym2) is not highly expressed in the naïve lung. However, both *Chil1* (BRP-39), encoded on chromosome 1, and *Chia* (AMCase) are detectable in naïve lung tissue^11^. Notably *Chia* is located immediately upstream of *Chil3* and would be susceptible to alterations from the targeting of *Chil3*. There was no difference in expression of *Chil1* or *Chia1* between wildtype, heterozygote, or knockout animals (**Fig. 6m and n**). Here, we have shown CRISPR-mediated deletion of Ym1 in this strain with no effect on the expression of the other associated chitinases or CLPs in the steady state. Considering how little is known about the function of Ym1, this strain will facilitate novel understanding of how this widely expressed protein functions in health and disease.

## Discussion

The C57BL/6 inbred strain of mice was originally established in the 1920s and has been well characterised as a model in many areas of research including cancer and immunology^39^. The C57BL/6 mouse was also selected as the reference genome for laboratory mice in 1999 by the Mouse Genome Sequencing Consortium (MGSC), the second mammalian genome sequenced after humans^40^. C57BL/6 was also selected as the background strain for the Knockout Mouse Project^41^, and assisted reproduction techniques are well established, facilitating the direct culture and genetic manipulation of C57BL/6 embryos. However, the strain is not without its limitations. Generations of inbreeding has inevitably led to genetic drift and strain differences between laboratories and suppliers. Notably, genetic duplications have been identified, including a copy number variant that disrupts the function of *Dock2* with implications on immune regulation^42^. Other genetic loss of function variants identified in C57BL6 include *Nnt*^43^, *Mnrn1*^44^, and *Rd8*^45^. In this study we attempted to create a conditional *Chil3* mouse model on the C57BL/6 background through CRISPR-Cas9. However, co-segregation of different gene edits, followed by ddPCR based copy number analysis, indicated a duplication of this locus in the supplied mice, aligning with previous studies which indicated a duplication of the *Chil3* locus in C57BL/6 strains^34,35^.

The BALB/c strain is an alternative mouse model for immunological research. After confirming a single copy of *Chil3* gene in BALB/c mice, we reasoned this would serve as a suitable background for generating a *Chil3*-deficient strain. We devised a whole gene deletion strategy exploiting the non-conserved sequences flanking the gene. However, we found BALB/c embryos difficult to generate through superovulation and natural matings. An experience also shared by several respondents to discussion on the International Society for Transgenic Technologies (ISTT) forum. Instead, we used embryos generated from F1 hybrids of the two strains. We reasoned that we would have a targetable single copy BALB/c allele in embryos, while potentially retaining the manipulation compatibility of C57BL/6 embryos. Whilst we were able to create a *Chil3* knockout allele using the CB6 strategy, the birth rates from embryos injected were still very low (1.7%). However, we should avoid definitive conclusions given the underpowered nature of this experiment. Recently alternative strategies have emerged that may allow the genetic manipulation of embryos *in situ*^46^. This would circumvent Using CB6 as a background strain does have the advantage of allowing more rapid backcrossing to each independent strain. In this study we backcrossed the *Chil3*-deletion strain with the BALB/c^olahsd^ strain. This backcrossing was accelerated using a speed congenic system. Using a panel of SNP markers, the contribution of specific genetic backgrounds was assessed allowing determination the optimal offspring to use for each crossing. This approach also allowed assessment of a suitable endpoint to confirm the animal was sufficiently backcrossed. Here we have reported data from just the BALB/c backcrossing, but in parallel we have also established the *Chil3* knock out allele on a backcrossed C57BL/6 background.

We have demonstrated the presence a recent duplication of the *Chia, Chil3*, and *Chil4* genes in C57BL/6J mice which is not present in BALB/c animals. This aligns with previous complete genome hybridisation data suggesting that this region had been duplicated in C57BL/6J mice^34^. Other studies have also shown that it is duplicated in C57BL/6N but not BALB/c animals^35^. Gene duplication events are considered to be one of the major drivers of genetic evolution^47^, allowing for the development of new gene functions from existing genetic code. Analysis of the the technically difficult harvesting and culture of embryos in challenging strains such as BALB/c.

protein expression in strains which harbour these duplications has shown that they correlate with increased protein expression in the serum. This demonstrates a direct functional consequence of these duplication events *in vivo*^35^. Understanding the specific functions of these duplicated genes, and differences between them will be key in determining their function in health and disease. Related to this study, we have also created a *Chil4* “knockdown” mouse on the C57BL/6 background^48^. In this strain we found that we had deleted one copy of the *Chil4* gene. This corresponded with a loss of secreted Ym2 into the airways during allergic airway inflammation, but Ym2 expression within epithelial cells was maintained. This again suggested variability between the two different copies in terms mechanisms of expression. Inbred mouse strains are common in research, yet widespread genetic difference between substrains and also within strains are becoming apparent. It is critical specific substrains of mice, and suppliers, that are used for experiments are well documented, in line with recent Laboratory Animal Genetic Reporting (LAG-R) framework^49^. This is vital to interpret biological results accurately but also generate robust tools for scientific discovery.

## Sup Figure Legends

**Sup Fig 4**: The presence of specific SNPs in the region described in Figure 4 allows for assessment of which CLP locus had been targeted. **a**) Schematic showing the possibly results of the identified SNP for genotyping the F1 mice their correlation with the allele which had been deleted.

## Additional Data

**Table 1:** Embryo survival rates.

| Strain | Total Injected | Embryo Survived (survival rate %) | Viable Pregnancy | Pups born (% of embryos injected) | Gene edited mice |
| --- | --- | --- | --- | --- | --- |
| BALB/c | 346 | 222 (64) | 1/10 | 3 (0.9) | 0 |
| CB6 | 177 | 144 (81) | 1/6 | 3 (1.7) | 1 |

## Materials and Methods

### Animals

BALB/c^OlaHsd^ and C57BL/6J^OlaHsd^ mice were purchased from a commercial supplier (Envigo, Hillcrest, UK), housed in individually ventilated cages within specific pathogen-free conditions at the University of Manchester Biological Services Facility. All animal procedures were performed in accordance with the UK Animals (Scientific Procedures) Act of 1986 under Project License (70/8548 and P4115856) granted by the UK Home Office and approved by the University of Manchester Animal Welfare and Ethical Review Body. Euthanasia was performed by asphyxiation in a rising concentration of CO_2_ followed by confirmation of death by cessation of circulation or cervical dislocation.

**Table 2:** Mouse strains.

| Item | Code |
| --- | --- |
| BALB/c<sup>OlaHsd</sup> | MGI:3047778 |
| C57BL/6J<sup>OlaHsd</sup> | MGI:2164189 |

### CRISPR-Cas9 Reagents

For the attempted conditional allele, we designed sgRNA specific to the introns flanking exon 3 for Chil3 that had low off target potential using https://wge.stemcell.sanger.ac.uk/. The 5’ sgRNA *ctgatattcttattctaccc* and 3’ sgRNA *atatggggtcgtataacatg* were subcloned into pUC57-sgRNA expression vector (Addgene 51132^50^). Vectors were linearised and used as a template for sgRNA synthesis using HiScribe (NEB) and RNA purified using Ambion MEGAclear transcription clean up kit, all according to manufacturer’s instructions. A dsDNA repair template comprising homology to the region flanking LoxP-exon3-loxP sequences was cloned.

For Chil3 gene deletion we designed two sgRNA targeting intergenic regions up- and down-stream of the gene (upstream sgRNA *attcccaaatctttaggaga, tactgtccacctcgagtggt*, downstream And *actgtccacctcgagtggtg*), and ssODN repair template *tgccatctctggggacacacagtggccttccacaggagattagattcccaaatctttaggactcgaggtggacagtacaatttcaccctctccactgtagagcacttcatcagagggc ta* to homologously repair across excised DNA. These reagents were all purchased as Alt-R products (Integrated DNA Technologies, Coralville, USA).

All editing reagents were purified or resuspended in sterile, RNase free Injection buffer (TrisHCl 1mM, pH 7.5, EDTA 0.1mM). For microinjection sgRNA were used at 20 ng/ml, EnGen Cas9 protein (NEB) 20 ng/ml and repair templates 10 ng/ml and 50 ng/ml respectively for dsDNA or ssODN DNA. Injection mixes were microinjected into one-day single cell mouse embryos from respective strains, generated through overnight matings following superovulation^51,52^. Zygotes were cultured embryos surgically implanted into the oviduct of day 0.5 post-coitum pseudopregnant mice. After birth and weaning, genomic DNA was extracted using Sigma REDExtract-N-Amp Tissue PCR kit and used to genotype pups. For the LoxP insertions we used flanking PCRs and sequencing to identify potentially positive pups, followed by re-amplification of products with high fidelity KOD polymerase, and subsequent blunt cloning into pCR-Blunt (Invitrogen). 12 colonies of transformed and miniprepped plasmid were Sanger sequenced using the M13R primer, and three different reads detected as indicated in figure 2d-e. We also integrated unique primer sequences in addition to LoxP sites to use for genotyping. Primer sequences are below, and position annotated in figure 2. For the gene deletion Primers immediately up and down stream of the sgRNA we used in combination. On an unedited allele no product would be generated, on a deleted allele a 384bp product is created, and was sequence confirmed. Primer sequences are below, and position annotated in figure 4. HDR mediated gene deletion was confirmed by Sanger sequencing.

**Table 3:**
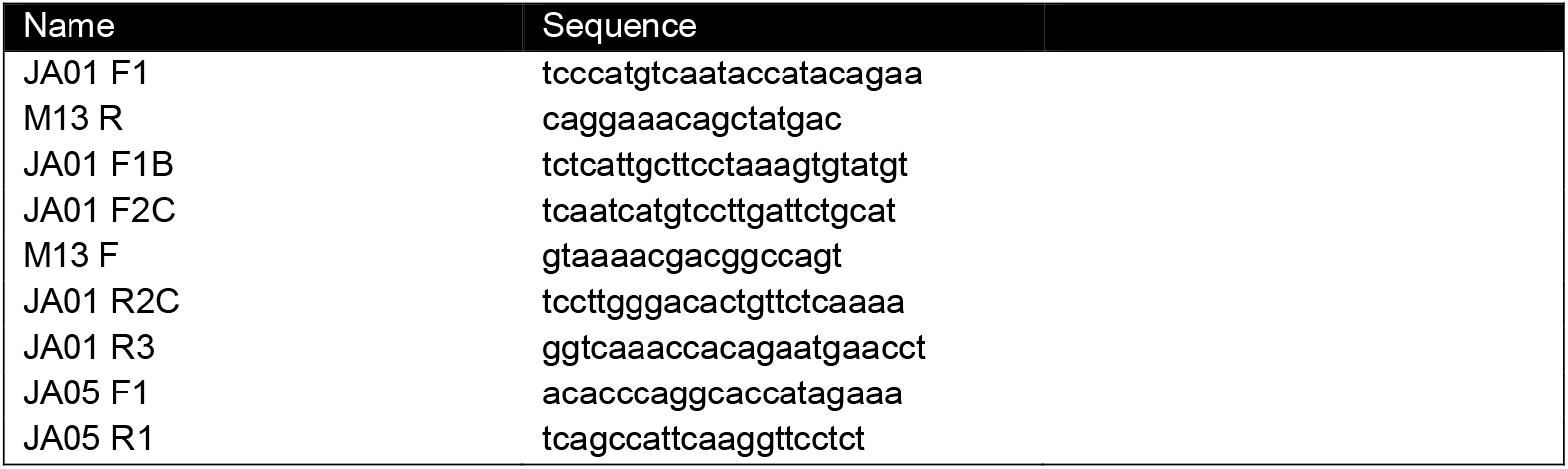
Primers used for genotyping and sequencing edited pups.

### Droplet Digital PCR

We used Droplet digital PCR (ddPCR) to confirm duplication in our mouse strains. ddPCR was conducted using the QX200 Droplet Digital PCR System (BioRad). Primers and probes were generated to respective CLPs (Table S1). Samples were generated as recommended by the manufacturer using the ddPCR Supermix for Probes (No dUTP) (186-3023), 900mM forward and reverse primers alongside 250mM hydrolysis probes. Oil emulsion was generated using the QX200 Droplet Generator and Droplet Generation Oil for Probes (1863005), DG8 Gaskets (1863009) and DG8 Cartridges (1864008).

Oil emulsions were then loaded onto ddPCR™ 96-Well Plates (12001925) and sealed with a PCR Plate Foil Heat Seal (1814045) using a PX1 PCR Plate Sealer (1814000). PCR products were then amplified using a C1000 Touch Thermalcycler (BioRad) using manufacturer suggested settings and an annealing temperature of 60°C (**Table 4**). Amplified droplets were then measured using the QX200 Droplet Reader and results analysed using QuantaSoftTM Analysis Pro to determine the copy number of the specific genes of interest in relation to a control gene *Tfrc* (Transferrin Receptor).

**Table 4:**
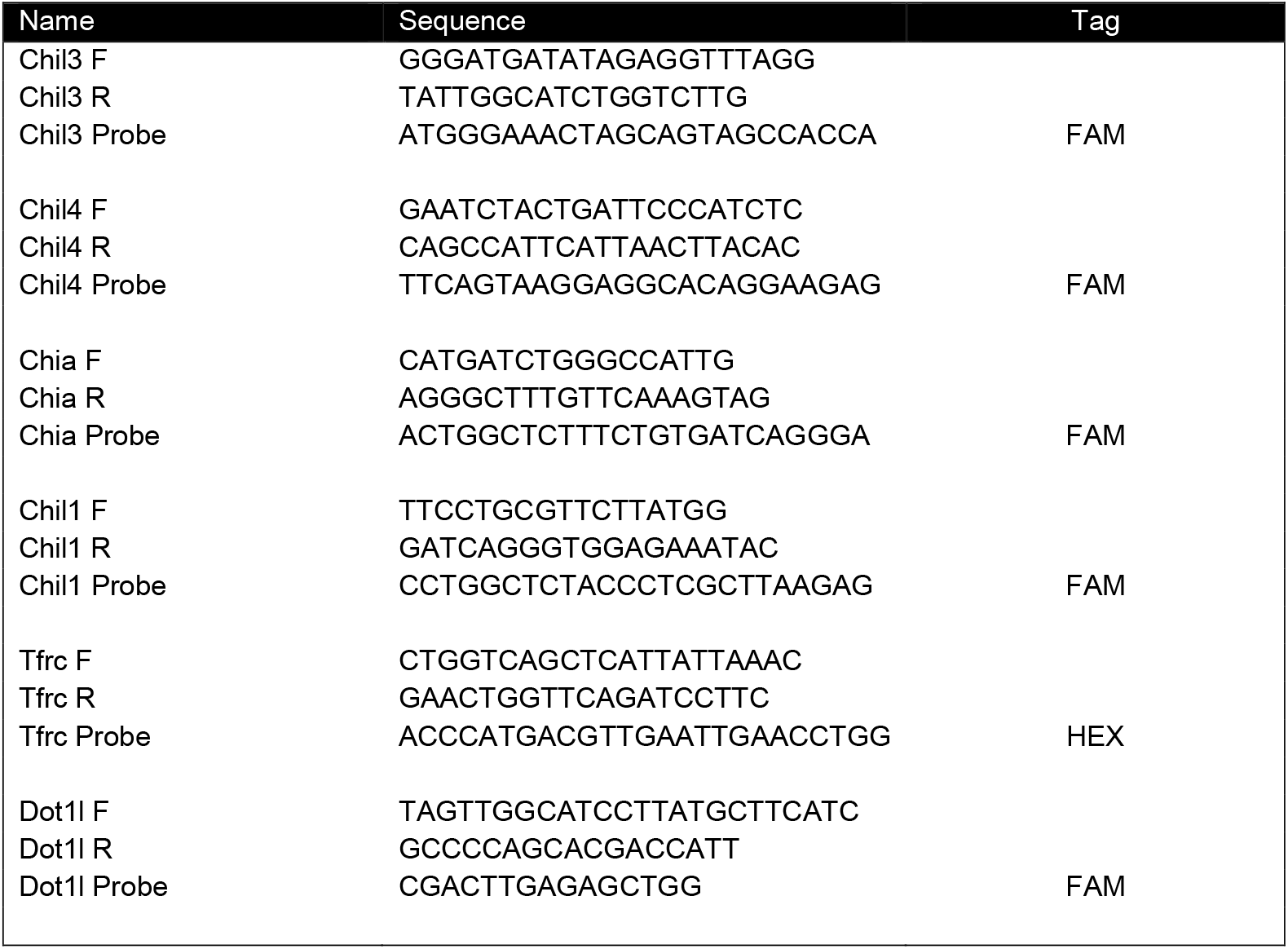
ddPCR primers and probes.

### Immunofluorescence

The left lung lobe was placed in 10% neutral-buffered formalin (NBF) (Sigma-Aldrich) for 24 hours before being transferred for storage in 70% ethanol (Fisher Scientific). Subsequently lungs were processed using an alcohol series and paraffin embedded. Sections of 5μm were cut using a microtome and mounted on slides (Superfrost Plus Adhesion). For immunofluorescence staining of lung tissue, sections were de-waxed through xylene and rehydrated through a series of alcohols before heat-induced antigen retrieval using 10 mM sodium citrate buffer, at pH 6.0. Samples were then blocked for nonspecific binding using 10% donkey serum in PBS (Sigma-Aldrich) followed by avidin biotin blocking (BioLegend). Primary antibodies were added to samples (**Table 5**) and incubated overnight at 4°C. Slides were washed in PBS before application of secondary antibodies (**Table 5**) for incubation at room temperature for 2 hours. Sections were mounted using coverslips and Fluormount G (Southern Biotech) containing 4’,6-diamidino-2-phenylindole (DAPI). Images of staining were captured using an EVOS FL imaging system (Thermo Fisher Scientific). To calculate Ym1 staining intensity, regions of interest (ROIs) were drawn around airways and embedded into the images using ImageJ software (version 1.51s). An ImageJ macro was used, with manual background threshold determined by a negative region, to calculate the integrated staining density across each ROI. Data was exported for graphing in GraphPad Prism (version 10.2.3).

**Table 5:** Antibodies used for immunofluorescence staining.

| Antigen | Antibody clone | Dilution | Manufacturer |
| --- | --- | --- | --- |
| **Ym1** | Goat polyclonal – biotinylated | 1:100 | R&D (BAF2446) |
| **Ym2** | Rabbit polyclonal | 1:1000 | Generated in-house |
| **Goat IgG** | Donkey anti-rabbit IgG 637 | 1:200 | R&D (NL005) |
| **Biotin** | Streptavidin 557 | 1:800 | R&D (NL999) |

### Cell Preparation

Bronchoalveolar lavage (BAL) was collected via cannulation of the trachea and washing of the airways four times with 0.4 mL PBS (Sigma-Aldrich). Samples were centrifuged (400 x g) and supernatant collected and stored at -80 °C. Superior and inferior lung lobes from the right side of the mouse were removed and chopped finely using blunt nosed scissors prior to the addition of digestion mixture containing 0.2 U/mL Liberase TL (Sigma-Aldrich) and 80 U/mL DNase Type 1 (Invitrogen) in pre-warmed RPMI (Gibco). Digestion was carried out for 45 minutes in a shaking incubator (220 rpm) at 37 °C. Digested tissue was mashed using a 70μm cell strainer (Greiner Bio-One) and red blood cells were lysed using ammonium-potassium-chloride (ACK) lysis buffer (Invitrogen). Single cell suspensions of digested lung tissue were then counted to acquire total live cell numbers using ViaStain AOPI staining solution (Nexcelom Bioscience) and a Cellometer Auto 2000 cell counter (Nexcelom Bioscience) before plating cells to stain for flow cytometry.

### Flow Cytometry

For surface staining, approximately equal numbers of cells per lung sample were washed in PBS prior to staining with Live/Dead (1:1000) (Thermo Fisher Scientific) and incubated with Fc-block (1:200) (CD16/CD32; clone 2.4G2; BD Biosciences) for 15 minutes. Antibodies (**Table 6**) were added in PBS containing 2mM EDTA and 1% FBS. For analysis of cytokines cells were stimulated for 4 hours at 37 °C with a cell stimulation cocktail containing protein transport inhibitor (1:500) (eBioscience). For intracellular staining, cells were fixed with fixation/permeabilization buffer (Invitrogen) before addition of permeabilisation buffer (Invitrogen). Cells were acquired on an LSRFortessa flow cytometer (BD Biosciences) using FACSDiva software (BD Biosciences) and analysed using FlowJo v10 software (BD Biosciences).

**Table 6:** Antibodies used in Flow cytometry.

| Antigen | Fluorophore | Clone | Dilution | Manufacturer |
| --- | --- | --- | --- | --- |
| **CD11c** | APC-Cy7 | N418 | 1:1000 | BioLegend |
| **CD11b** | BV711 | M1/70 | 1:1000 | BioLegend |
| **CD45** | BV785 | 30-F11 | 1:400 | BioLegend |
| **CD64** | BV605 | X54-5/7.1 | 1:400 | BioLegend |
| **Siglec-F** | BV421 | E50-2440 | 1:400 | BD Biosciences |
| **Ly6G** | AF647 | 1A8 | 1:400 | BioLegend |
| **MHCII (I-A/I-E)** | BV650 | M5/114.15.2 | 1:1000 | BioLegend |
| **Ym1\*** | PE-Cy7 | Polyclonal | 1:200 | Novus |

\*Biotinylated Ym1 used at 1:200 and detected using streptavidin (1:400)

### qRT-PCR

The post-caval lung lobe was collected in RNA Later solution (Thermo Fisher Scientific) and stored at -80 °C before RNA extraction using the PureLink RNA Mini kit (Invitrogen) according to manufacturer’s instructions. RNA yield and purity were determined using a NanoDrop™. RNA (500 ng) was reverse transcribed using 5x RT buffer (Bioline), Tetro reverse transcriptase (Meridian Bioscience), oligo dT 15-mer (Integrated DNA Technologies), Ribonuclease inhibitor (RNasin) and 10 mM deoxynucleoside triphosphates (Promega). Quantitative real-time PCR was performed using SYBR green master mix (Agilent Technologies), specific primer sets (**Table 7**, Integrated DNA Technologies), and read on a LightCycler 480 II (Roche). Changes in gene expression levels were determined using the ΔΔCq method relative to the geometric mean of housekeeping genes *RpI13a* and *Rn18s*.

**Table 7:**
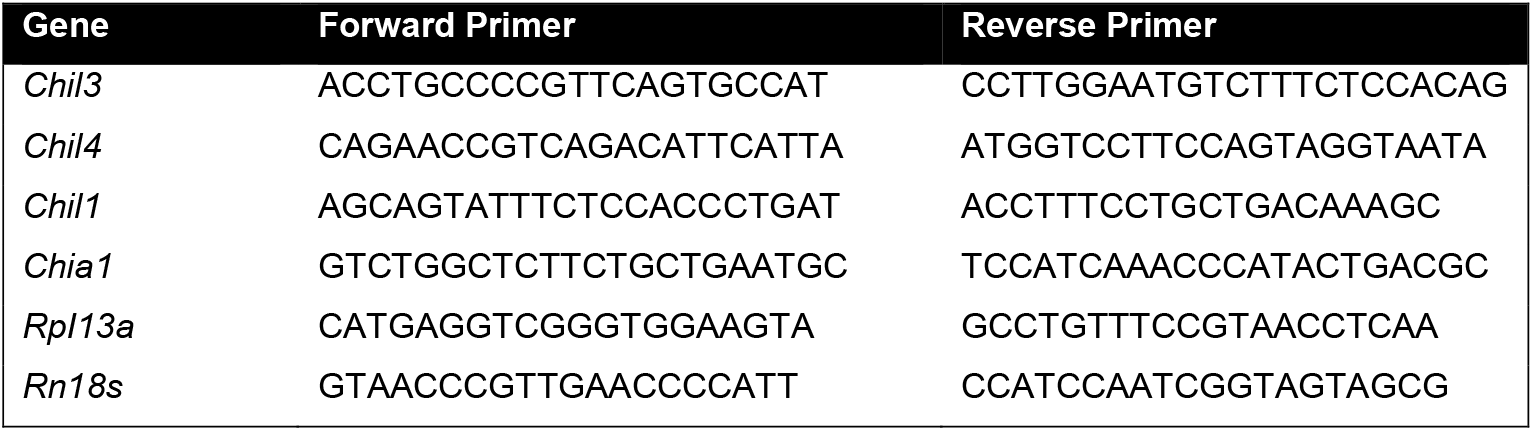
Primer sequences for detection of mRNA of listed genes by quantitative reverse transcription PCR.

### Statistical Analysis

GraphPad Prism (version 10.2.3) was used for statistical analysis. Normality of data was assessed using the Shapiro-Wilk test with some datasets log2 transformed to achieve normal distribution. For normally distributed data parametric tests were used and for non-normally distributed data, non-parametric tests were used. For comparisons between multiple groups, one-way analysis of variance (parametric) with Holm-Sidak’s multiple comparisons test or Kruskal-Wallis test (non-parametric) with Dunn’s multiple comparison test were used, as indicated in figure legends. Data are represented as mean± SD and exact P values are shown on graphs.

## Acknowledgements

We thank Brian Chan and Adam Johnson for technical support as well as the flow cytometry, bioimaging, and genomic editing unit core facility at the University of Manchester.

## Contributions

Conceptualization: JEP, JEA, ADA, and TES

Methodology: JEP, NEH, and ADA

Formal analysis: JEP and GEB

Investigation: JEP and GEB

Writing - Original Draft: JEP, GEB, and ADA

Writing - Review & Editing: JEP, GEB, PHP, JEA, ADA, and TES

Visualization: JEP, ADA, and GEB Supervision: ADA, JEA, PHP, and TES Funding acquisition: ADA, JEA, and TES

