## Extended Figure 4 for "Generation of a Ym1 Deficient Mouse utilising CRISPR-Cas9 in CB6 Embryos"

a

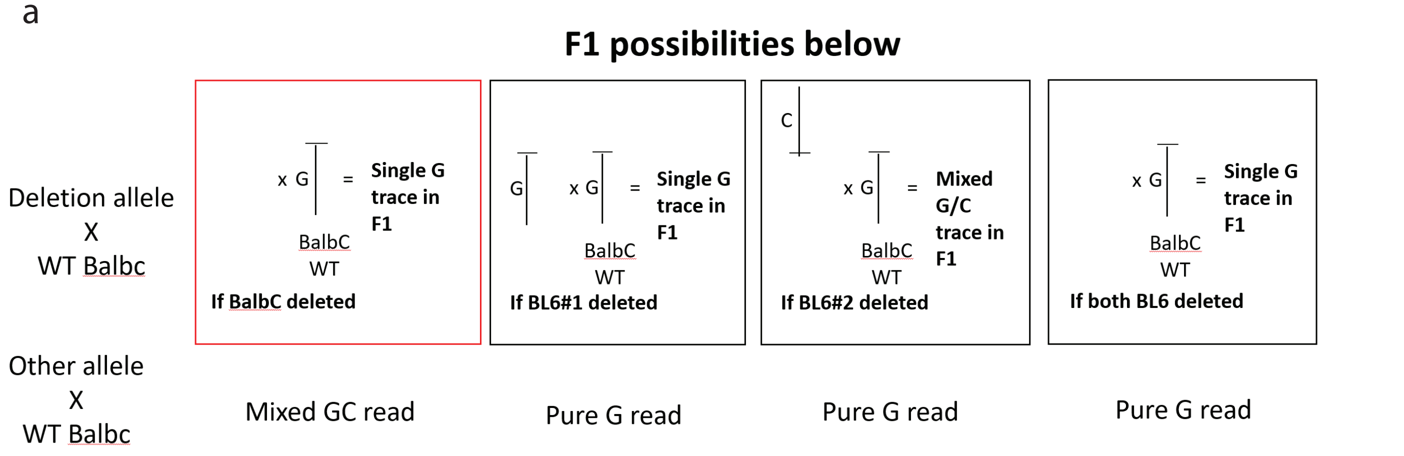


So if we get a single G trace in the PCR positive, and a mixed GC read in the PCR negative it could be what we want
